# Human endogenous retrovirus K (HERV-K) envelope structures in pre- and post-fusion by Cryo-EM

**DOI:** 10.1101/2025.04.04.647320

**Authors:** Jeremy Shek, Chen Sun, Elise M. Wilson, Fatemeh Moadab, Kathryn M. Hastie, Roshan R. Rajamanickam, Patrick J. Penalosa, Stephanie S. Harkins, Diptiben Parekh, Chitra Hariharan, Dawid S. Zyla, Cassandra Yu, Kelly C.L. Shaffer, Victoria I. Lewis, Ruben Diaz Avalos, Tomas Mustelin, Erica Ollmann Saphire

## Abstract

The HERV-K envelope glycoprotein (Env) is aberrantly expressed in diseases including cancers and autoimmune disorders and is targeted by antibodies. The lack of structural information has hindered functional and immune recognition studies. We solved structures of the HERV-K Env in both pre- and post-fusion states with novel monoclonal antibodies using cryo-EM. The pre-fusion Env assembles as a trimer with a distinct fold and architecture compared to other retroviruses, while the post-fusion conformation features a unique “tether” helix within the TM subunit. A panel of monoclonal antibodies, elicited to facilitate structure determination, have been characterized for conformational and subunit specificity, serving as valuable research tools. These findings establish a structural framework for mechanistic studies of HERV-K Env in diseases and evaluation as a potential therapeutic target.

## Introduction

Over millions of years, retroviruses have infected humans and our primate ancestors, integrating into our DNA and passing down through generations. Records of these ancient viral infections are now scattered throughout the human genome, known as human endogenous retroviruses (HERVs). Genomic analyses have revealed that HERV sequences comprise 8% of the human genome, with approximately 98,000 annotated insertions(*1*). HERVs possess a canonical retroviral genome like their exogenous counterparts with *gag*, *pol*, and *env* genes flanked by long-terminal repeats (LTRs) containing regulatory elements(*2*, *3*). Over time, most HERV elements accumulated mutations or deletions that rendered them non-functional and incapable of producing infectious virions(*3*).

Among these HERVs, the HERV-K (HML-2) family is the most recently acquired insertion; HERV-K insertions have accumulated the least number of deleterious mutations and are the most transcriptionally active. There are ∼100 HERV-K full proviral inserts in our genomes, many of which contain complete open reading frames (ORFs) for each of the viral proteins(*4–6*). While epigenetic modifications and transcriptional repressors such as CpG methylation, histone acetylation, and chromatin remodeling keep HERV-K transcriptionally silenced in healthy adult cells(*7*, *8*), recent studies reveal that in certain disease states, HERV-K proteins escape suppression and are actively transcribed(*9–11*).

Specifically, expression of the HERV-K envelope glycoprotein (Env) protein, encoded by the *env* gene, has been detected in breast, ovarian, prostate, melanoma, leukemia, and other cancers(*12–21*). Aberrant Env expression is associated with cancer cell proliferation, tumorigenesis, and disease progression(*22–27*). Autoimmune diseases including rheumatoid arthritis (RA), systemic lupus erythematosus (SLE), and type I diabetes (T1D) have also been linked with increased expression of Env(*5*, *28–33*). Furthermore, it is evident that the immune system develops an antibody response against Env in many cases and that antibodies against Env may even contribute to the pathogenesis of these malignancies(*5*, *13*, *31*, *34–37*). Although much remains to be discovered about roles of Env in disease, the aberrant expression of Env on the cell surface could be leveraged for immunotherapy and diagnostics. Indeed, antibodies targeting HERV-K Env have demonstrated antitumor effects against breast cancer(*38*), engineered chimeric antigen receptor (CAR) T cells targeting Env can inhibit tumor metastasis(*39*, *40*), and mice immunized with a vaccine directed against Env can reject tumors expressing HERV-K Env(*41*). Further, antibodies targeting HERV-K Env neutralize Env-mediated neurotoxicity in the cerebral spinal fluid of amyotrophic lateral sclerosis patients (ALS) by blocking binding of Env to its receptor(*33*, *37*, *38*). Collectively, these findings suggest Env as a marker or direct therapeutic target in a range of diseases.

Confounding molecular analysis of HERV-K Env function, recognition, and targeting is the lack of any structural information of the HERV-K antigen. While every human bears over ten complete copies of HERV-K *env* with the capacity for expression in their genomes, we do not yet know what the protein looks like or how antibodies may target it. In this study, we use stabilizing approaches, antibody discovery, and cryogenic electron microscopy (cryo-EM) to reveal the first structures of the HERV-K envelope glycoprotein ectodomain in both the pre-fusion and post-fusion conformations. Our findings report an overall architecture that is distinct from all known retroviral structures. We also illuminate surfaces available for immune recognition and monoclonal antibodies that begin to chart the antigenic landscape of HERV-K Env. The stabilized protein and mAbs open doors for functional studies of Env in disease and structure-based development of new interventions.

## Results

### Anti-HERV-K Env monoclonal antibody discovery

The lack of research reagents for studying HERV-K Env and its subunits hindered our initial study of the protein. The commercially available monoclonal antibodies (mAbs) HERM 1811-5 and HERM 1821-5 (Austral Biologicals) were elicited by immunization with *E. coli-*produced glycoproteins. They recognize linear epitopes and therefore cannot discern properly folded from misfolded protein and were not suitable for structural studies of the HERV-K Env.

In this study, we immunized BALB/c mice with the wild-type Env ectodomain (Env_Ecto_) produced in *Drosophila* S2 insect cells. Using the Bruker Beacon antibody discovery platform, followed by biochemical characterization, we selected a subset of ten monoclonal antibodies from the panel that would together broadly cover the range of conformational (pre- and post-fusion) and subunit (SU and TM) specificities (fig. 1a). We screened antibodies on ELISA coated with either purified wild-type (WT) unstabilized Env_Ecto_, which contains both SU and TM_Ecto_ in a mixture of pre- and post-fusion conformations, or with post-fusion TM_Ecto_ alone. Six of the mAbs are bound to both WT Env_Ecto_ and post-fusion TM_Ecto_ while the remaining four are only bound to WT Env_Ecto_ (fig. S1a). To determine whether these mAbs recognize linear or conformational epitopes, we denatured WT Env_Ecto_ or post-fusion TM_Ecto_ at 95°C with DTT for 10 minutes and screened the mAbs using ELISA again. Six mAbs bound to either denatured Env_Ecto_ or post-fusion TM_Ecto_, revealing that they likely bind to linearized epitopes. Six mAbs bound to either denatured Env_Ecto_ or post-fusion TM_Ecto_, revealing that they likely bind to linearized epitopes. Kenv-6 only binds WT Env, and fails to bind denatured or postfusion Env_Ecto_, indicating it is pre-fusion specific and recognizes a conformational epitope. In contrast, while Kenv-4 binds more strongly to post- fusion TM_Ecto_ (fig. S1a). Western blot analysis using WT Env_Ecto_ was performed with each mAb to further confirm subunit and conformational specificities (fig. S1b). Two mAbs, Kenv-2 and Kenv-3, did not bind to fully denatured protein but did recognize Env in western blots.

**Figure 1.**
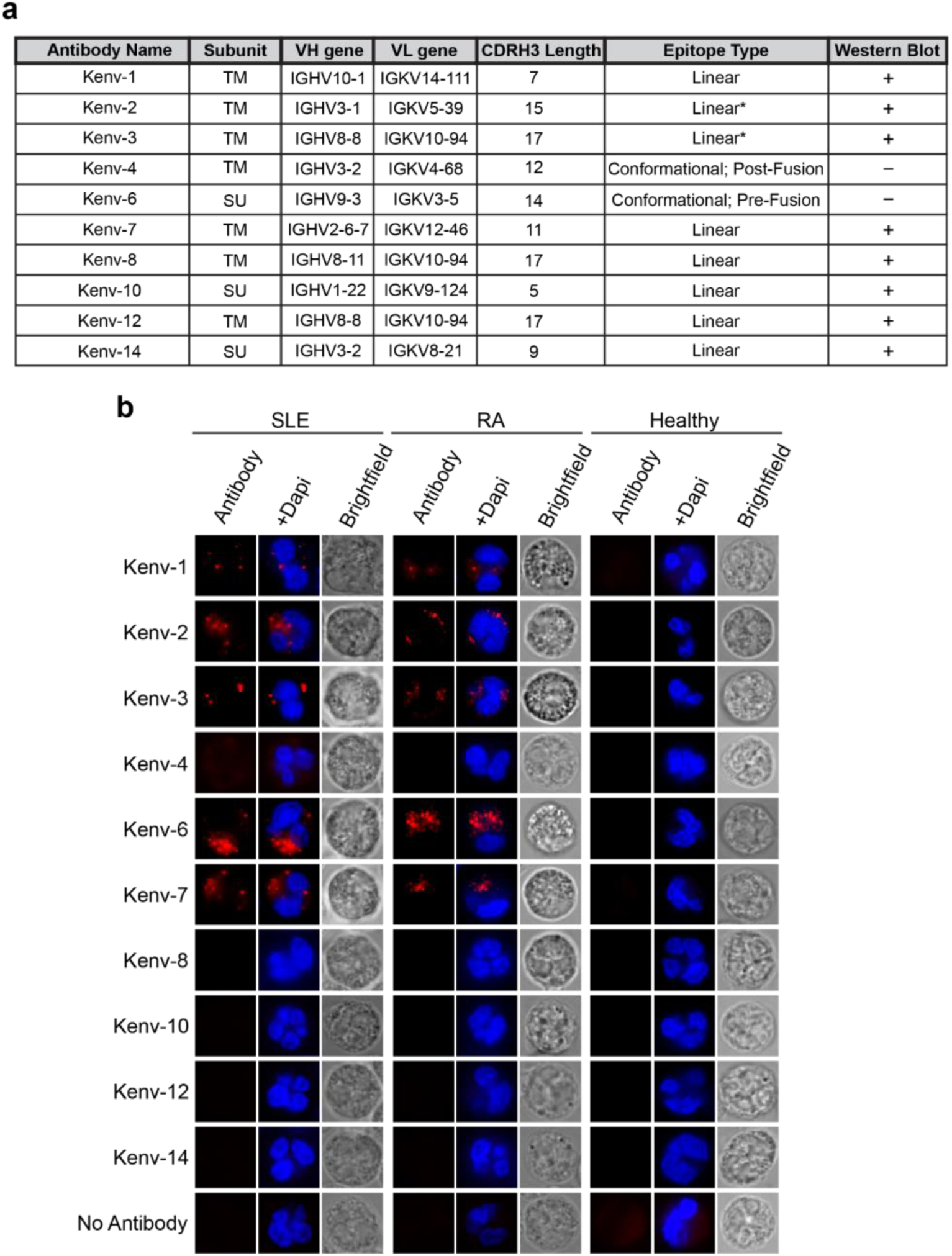
Novel monoclonal antibodies against HERV-K Env. **a.** Table of anti-Env mAbs discovered in this study. Antibodies were screened for surface protein (SU) or transmembrane protein subunit (TM) binding using ELISA shown in fig. S1a. Epitope type was determined using ELISAs in combination with western blots (fig. S1b). * = presumed linear based on western blotting results. **b.** Immunofluorescence staining of isolated neutrophils from either SLE, RA, or healthy donors using mAbs from this study. Neutrophils from RA samples were treated with IFNα prior to fixation. Healthy neutrophils isolated with or without IFNα treatment show no staining by any mAb.

Together, we found that three mAbs are SU specific (Kenv-6, Kenv-10, Kenv-14) and seven are TM specific (Kenv-1, Kenv-2, Kenv-3, Kenv-4, Kenv-7, Kenv-12). Two antibodies, Kenv-6 and Kenv-4, bind to conformational epitopes on the properly folded SU and post-fusion TM, respectively (Figure 1a). With these novel monoclonal antibodies, we could reliably detect each Env subunit when engineering Env to stabilize the pre-fusion conformation. Further, the two conformational mAbs facilitated the structure determination of Env in pre- and post-fusion states.

We sought to further validate antibody recognition to see if these mAbs would recognize HERV-K Env expressed in human patient samples as well as recombinant protein. We found that five of these antibodies, Kenv-1, Kenv-2, Kenv-3, Kenv-6, and Kenv-7 are able to stain isolated neutrophils from both SLE and RA patients (fig. 1b), which have been previously reported to express Envs from two HERV-K loci: HERV-K102 and K108 (*5*, *30*, *31*, *35*). In contrast, these mAbs did not have any level of detectable antibody binding to neutrophils from healthy patients.

### HERV-K Env Pre-fusion Trimer Stabilization

There are multiple insertions of the human endogenous retrovirus K (HERV-K) in the human genome. Among them, at least 10 loci contain intact open reading frames for the *env* gene(*5*). These HERV-K family *env* genes are classified by the absence (type 1) or presence (type 2) of a 292-bp sequence at the 5’ end of the gene, which encodes a signal peptide for plasma membrane surface localization(*42*). Previous studies have generated consensus sequences of the HERV-K proviral genome by comparing HERV-K proviral inserts with the type 2 *env*s. These consensus sequences generate infectious virions(*43*, *44*), are fusogenically active, and have broad tropism(*45*). In this study, we used the type 2 *Phoenix* consensus Env ectodomain (residues 97-632) sequence(*43*)due to >97% sequence identity to all Envs with the potential for expression (fig. S2a).

HERV-K Env is a class I viral fusion protein comprising an expected receptor-binding/surface protein subunit (SU) and a fusion/transmembrane protein subunit (TM). Three non-covalently bound SU-plus-TM protomers trimerize on the cell surface(*46*). Consistent with the metastability of class I viral fusion proteins, wild-type Env can spontaneously adopt the more energetically stable post-fusion conformation, in which the SU subunits disassociate, leaving the TM subunits to fold into a six-helix bundle (6HB)(*47*, *48*). This metastability poses a significant obstacle to the expression and stability of pre-fusion conformation Env protein that would be amenable to structure determination by cryo-EM.

To overcome the problems associated with glycoprotein metastability, we explored methods that have been previously successful in the engineering of other viral envelope glycoproteins, introducing functional mutations to confer stability to the pre-fusion conformation. Modifications we designed include (i) helix-breaking proline mutations, (ii) inter-protomer disulfide linkages, and (iii) additional multimerization domains(*49–51*). We screened and tested mutants for expression, post-translational processing, and oligomerization, employing an iterative process of AlphaFold- and DeepCoil-driven predictions(*52*, *53*) to determine where to place helix-breaking mutations to break the long heptad repeat 1 (HR1) helix that would otherwise drive formation of the post-fusion six-helix bundle. 22 mutants with one or two prolines within the HR1 sequence were evaluated for expression and oligomerization. The best-behaved sequences were then input into AlphaFold to generate new model predictions that were more “pre-fusion-like”. From these models, we next designed 13 potential cysteine pairs using structural visualization in ChimeraX(*54*, *55*) and the Disulfide by Design 2.0 server(*54*, *55*) to form disulfide bonds between SU and TM_Ecto_. In parallel, we assessed the utility of heterologous oligomerization domains and modified linker lengths, and evaluated expression and post-translational processing in different eukaryotic cell lines.

Among the hundreds of mutations and modifications we screened, the combination of one introduced cysteine pair between SU (V437C) and TM_ecto_ (V498C), two point mutations that altered the furin cleavage motif from RSKR to RRRR, and addition of the T4 Fibritin trimerization domain at the C terminus of TM_Ecto_ together resulted in the expression of stable, pre-fusion Env_Ecto_ trimers (fig. 2a, fig. S3). Size exclusion chromatography and SDS-PAGE analysis of Env_Ecto_ expressed in *Drosophila* S2 insect cells indicate that Env_Ecto_ is efficiently processed by furin into SU and TM_Ecto_ subunits which are joined by a disulfide bond, and that this Env oligomerizes as a stable trimer in solution (fig. S3b-e). We assessed these Env_Ecto_ constructs using nano differential scanning fluorimetry and demonstrated an increase in melting temperatures from 46.0°C in the unstabilized to 53.8°C in the stabilized Env (fig. S3f-h). Env_Ecto_ trimers produced and purified in this way were complexed with Fab fragments from the novel Kenv-6 mAb for cryo-EM. 479,625 particles from 13,141 movies were processed with CryoSPARC(*56*) to obtain a reconstruction at a 2.2 Å global resolution (fig. 2b).

**Figure 2.**
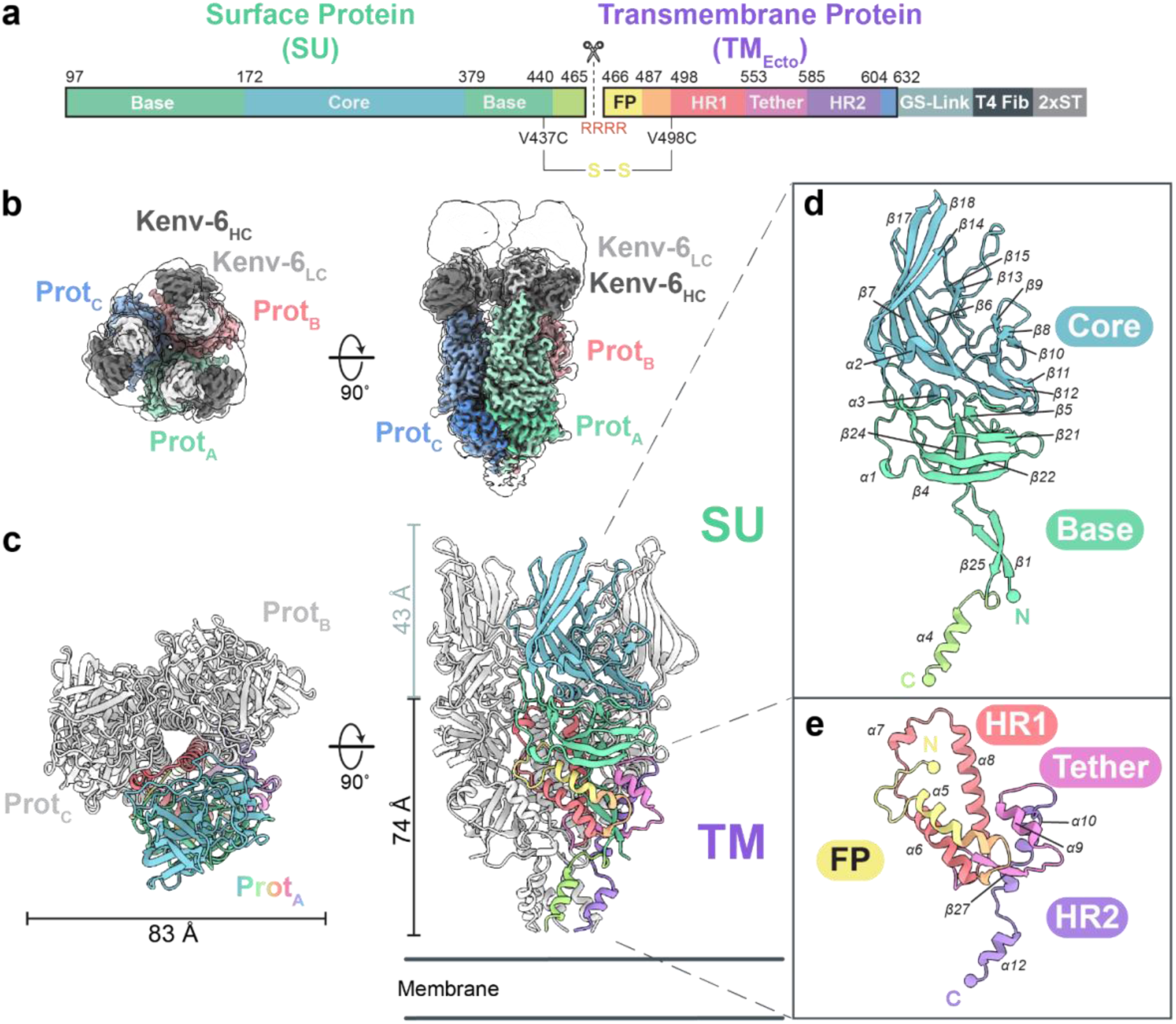
Cryo-EM structure of the pre-fusion HERV-K envelope glycoprotein trimer. **a.** Primary structure annotation of HERV-K Env_Ecto_ construct. Domains are color-coded. Abbreviations: FP, fusion peptide; HR1, heptad repeat 1; HR2, heptad repeat 2; GS-Link, 12-amino acid GGS linker; T4-Fib, T4 Fibritin foldon trimerization domain; 2xST, Twin Strep-Tag II. Regions boxed in black are sequences of Env_Ecto_. Black scissors indicate the furin cleavage site that cleaves Env into its SU and TM subunits. Engineered disulfide bond mutations and furin-cleavage motif modifications are annotated. **b.** 2.2 Å resolution cryo-EM map of HERV-K Env_Ecto_ trimer (colored using green, red, and blue shades) bound to three copies of the Kenv-6 Fab. Densities corresponding to the three SU subunits are colored with the lighter shade and densities for the three copies of TM are colored in the darker shade. A low-resolution map is overlaid and represented as a silhouette. **c.** Cartoon representation of the HERV-K Env_Ecto_ trimer. Protomer A is colored in a rainbow as described in (**a**). The remaining 2 protomers are colored in white. **d.** Isolated cartoon representation of the surface protein (SU) subunit from an individual protomer. **e.** Cartoon representation of the transmembrane protein ectodomain (TM_Ecto_). Both **d** and **e** are colored using the same color scheme as (**a**,**c**). N- and C-terminal Cα atoms for each subunit are represented as spheres.

### The architecture of the pre-fusion HERV-K Env_Ecto_ Trimer

With the high-resolution map we obtained, we were able to build nearly the entirety of Env_Ecto_ in the pre-fusion conformation with only residues at the N- and C-terminal ends of the SU (97–99, 460–465) and the N-terminal, membrane-proximal end of TM_Ecto_ (621–632) unmodeled (fig. 2b,c). Overall, the soluble HERV-K Env_Ecto_ pre-fusion trimer adopts an elongated shape akin to an inverted tripod with a height of ∼117 Å along the vertical axis and a width of ∼83 Å. In the trimer, the SU subunits sit above their TM counterparts and are shaped like prongs that extend 49 Å from the highest point of the TM_Ecto_ near the trimer axis (fig. 2c). At the trimer apex, between the assembled three SUs, is an open central cavity with a solvent-accessible surface area of approximately 7,000 Å. Though the furin cleavage motif (residues 461-465) is unmodeled, the C terminus of SU ending just before the furin motif at residue 459 folds as a downward-pointing alpha-helix (α4) toward the membrane surface, and forms a bundle with the C-terminal helices (α12) of the TM_Ecto_ (fig. 2c,d). The distance between the two residues flanking the furin sequence (V459 on the SU and F466 on TM) is 53 Å. If uncleaved, this seven amino-acid span could reach ∼25 Å. A separation of 53 Å indicates that upon cleavage, the newly formed termini must displace from each other, with the newly formed SU C terminus moving toward the foot of the trimer while the newly formed TM_Ecto_ N terminus burying itself toward the central coiled-coils. This ∼30 Å displacement explains why attempts to engineer an uncleavable Env_Ecto_ did not result in successful expression or trimerization, and instead resulted in misfolded proteins with improper quaternary structure.

#### Structure of the Surface Protein Subunit (SU)

The SU (residues 97-465) is a 369 amino-acid polypeptide comprising 4 α helices and 25 β strands. The SU consists of two main subdomains, here termed the core and the base (fig. 2d).

The core domain, formed by residues 172-378, is membrane-distal at the trimer apex, and is likely involved in receptor binding to heparin, CD98HC(*33*, *57*), and perhaps other factors yet to be discovered. The most prominent feature of the SU core is a β sandwich in which one side, comprised of five anti-parallel strands (β6, β7, β14, β17, and β18), forms an elongated, distorted sheet, while the opposing sheet is formed by two short anti-parallel strands (β13 and β15). Two disulfide bridges stabilize key loops in the core structure. The first, C275-C282, stabilizes a surface-exposed loop at the apex of the SU. This loop may be important for receptor engagement: an antibody targeting the adjacent β14 strand blocks the interaction of Env with CD98HC(*33*), an integrin-associated protein that mediates integrin-dependent signals that promote tumorigenesis(*58*). The second disulfide, C227-C246, anchors the loop flanked by the two cysteine residues, the largest and most flexible loop in our model. At higher map thresholds (e.g., 11σ), the volume for the residues 233-239 of this loop are unresolvable. Applying a blur factor of +50Å^2^ revealed continuous density, which allowed us to trace and model the loop in its entirety.

The SU base domain, formed by residues 97-171 plus 379-439, is the more membrane-proximal of the core-base pair and is responsible for mediating interactions with the TM subunit in the pre-fusion conformation. The base consists of a distorted β barrel structure (β2, β4, β5, β20, β21, β22, and β24) positioned above the β1-β25 sheet where the N- and C termini join. This β sheet extends downward toward the membrane. The C-terminal end of the SU is proximal to the very bottom of the ectodomain, where α4 is located, directly preceding the furin cleavage motif (residues 461-465), which is not resolved. The SU subunit does not form any contacts with neighboring SU subunits in the trimer in our static structure and does not appear to contribute to trimeric interactions.

#### Structure of the Pre-Fusion Transmembrane Protein Subunit (TM)

The transmembrane subunit (TM) is responsible for the membrane fusion mechanism of retroviral Env. The pre-fusion TM ectodomain structure presented here comprises eight α helices and two β strands (fig. 2e). Consistent with class I viral fusion proteins, TM_Ecto_ contains a hydrophobic fusion peptide followed by heptad repeat sequence motifs, HR1 and HR2. The fusion peptide (residues 466-486) begins as an unstructured coil that transitions into α5, sitting above α6 of HR1. The first heptad repeat motif, HR1 (498–552), is divided into three α helices: α6, α7, and α8. The third, α8, is amphipathic. The hydrophobic side of the α8 helix faces the center of the trimer axis and forms the central coil. HR1 and HR2 are linked by a 32-residue tether region that contains a CX_7_C disulfide motif (C551-C559) specific to betaretroviral envelope proteins (*46*). HR2 (residues 585-603) extends proximally toward the membrane, with the C-terminus of TM_Ecto_ forming a bundle at the bottom of the trimer (Fig 2c,e).

#### The Pre-Fusion TM Wraps Around SU Base

Within each protomer, the SU contacts its TM partner via the 7-stranded β barrel and the terminal β1-β25 sheet of the base domain. The pre-fusion TM_Ecto_ encircles the extended N- and C-terminal strands of the base, forming a C-shaped clasp (fig. 3b,c). Helices α6 and α7 partially envelop the extended N- and C-terminal strands of SU before reversing direction and extending behind where β27 hydrogen bonds to β25, joining the β1-β25 sheet in an anti-parallel fashion (fig. 3d). The TM completes its wrapping around the SU with α9 enclosing the remaining portion of the clamp. Sidechain nitrogen atoms from R576 and W572 form a hydrogen bond network with the backbone oxygens of H492 and S493, effectively closing and locking the clasp (fig. 3b). As a result, the HERV-K TM forms a near-complete ring underneath the SU base around the terminal β sheet (fig. 3c).

**Figure 3.**
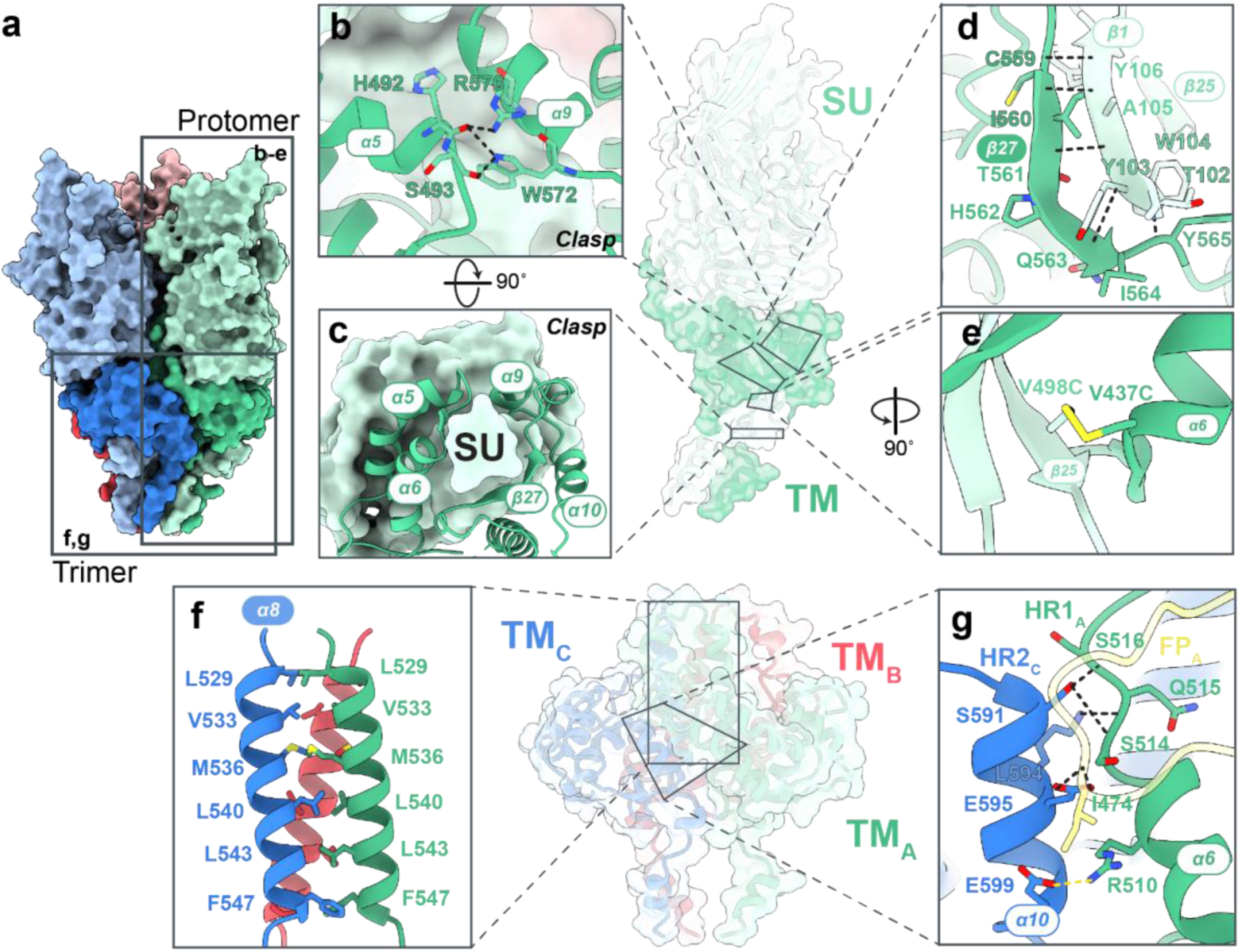
Pre-fusion Env_Ecto_ Interfaces. **a.** The pre-fusion Env_Ecto_ trimer displayed using a molecular surface representation. SU+TM protomer interactions are shown in panels (**b-e**) and trimeric interactions are shown in panels (**f,g**). **b.** The clasp formed by the TM around the base domain of the SU viewed from the front. Residues involved in clasp locking are shown as sticks. **c.** The C-shaped clasp formed by the TM around the SU viewed from bottom-up. **b,c.** The SU is shown as a light green surface and the TM is a dark green cartoon. **d.** β sheet formation between the β1 strand from the SU (light green) and the β27 strand from the TM (dark green). **e.** The engineered disulfide bond between the SU and TM to stabilize Env_Ecto_. **f.** Side-view of the central coiled-coil. Hydrophobic residues facing the interface from each helix are shown as sticks. **g.** The interactions between adjacent pre-fusion TM subunits distal from the central coils. The fusion peptide of TM_A_ is colored in yellow. **b,d,g.** Hydrogen bond interactions are shown by dashed black lines and salt-bridges are shown in dashed yellow lines.

Dissociation of the receptor-binding subunit from the fusion protein subunit is a trigger of conformational change to the post-fusion state(*47*, *48*). To prevent the dissociation of the non-covalently bound SU and TM subunits, part of our previously described engineering was the introduction of an intersubunit disulfide between residues 437 on SU and 498 on TM (fig. 3e). The engineered disulfide anchors the N terminus of the TM, between the fusion peptide and HR1 at the membrane-proximal end of strand β25, to the base of SU. This disulfide stabilizes the pre-fusion conformation of Env by preventing the HR1 refolding and extension that would be needed for transition to the post-fusion conformation (fig. S7b).

#### TM-trimer interaction

Much like other class I viral fusion proteins such as the HIV Env, Ebola GP, RSV F, and Influenza hemagglutinin (HA) proteins, the Env trimer is stabilized by the presence of a central coiled-coil(*47*, *59–62*). In HERV-K Env, the coiled-coil is formed by helix α8 (residues 529-548) in the HR1 sequence of each TM subunit, via hydrophobic interactions between the a and d positions of the heptad repeats at the central axis (fig. 4f). We found that both the length of the α8 helix as well as the *absence* of helix-breaking mutations are crucial for Env trimerization. We initially followed the HIV-1 SOSIP strategy(*63*) of inserting kinks within the junction between α7 and α8 of Env, mutating residues to prolines, and did notice an increase in expression with three such mutants. However, our best-expressing mutant, L529P, yields Env monomers alone. Even if fused to a T4 Fibritin trimerization domain, L529P Env remains monomeric and does not trimerize. We hypothesize that the proline insertion destabilizes the secondary structure of the N terminus of α8, decreasing the number of hydrophobic interactions in the central coiled-coil (fig. 4f), and thereby weakening the trimer complex. Trimerization of Env_Ecto_ was achieved by reverting P529 back to its native lysine.

**Figure 4.**
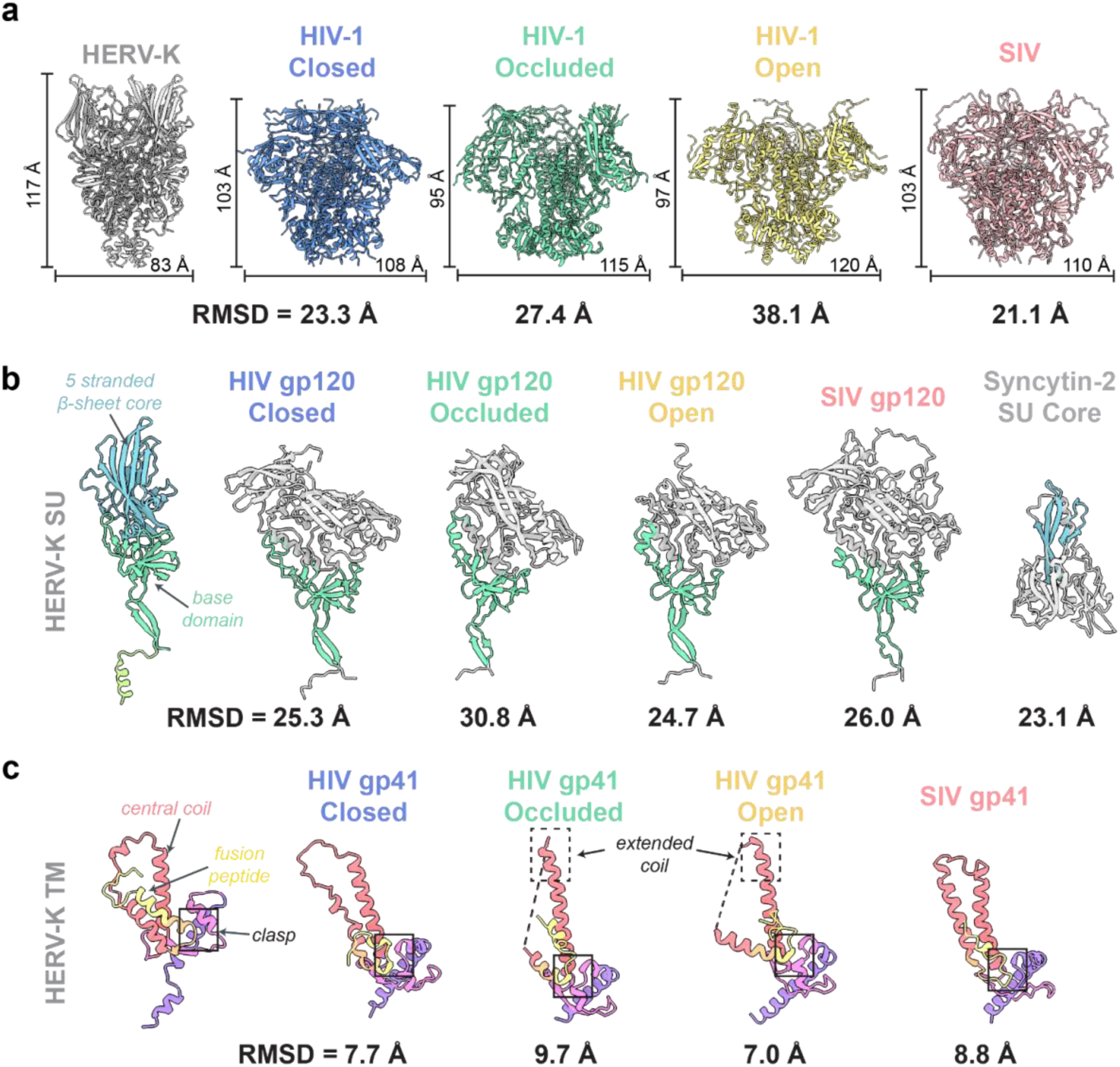
Comparison of pre-fusion HERV-K Env_Ecto_ to other solved pre-fusion retroviral structures. **a.** Models of HERV-K Env, HIV gp (closed (PDB:5CEZ), occluded (PDB:7TFO), open (PDB:5VN3), and SIV (PDB:7T4G) ectodomain trimers. Approximate heights and widths of each complex were measured using ChimeraX (*54*). **b.** Comparison of receptor-binding subunits (SU: HERV-K, Syncytin-2; gp120: HIV, SIV). HERV-K Env SU is colored by subdomain while the other receptor-binding subunits are colored in light gray. Shared structural features to HERV-K SU are colored and labeled with their respective colors. **c.** Comparison of fusion protein subunits (TM: HERV-K; gp41: HIV, SIV) in pre-fusion conformation. HERV-K Env TM is colored by subdomain. Similar structural features presented on the HIV and SIV gp41 subunits are colored similarly to TM. The C-shaped clasp that encloses the SU/gp120 base domain is boxed in black on each TM/gp41. The HR1 central coil in the occluded and closed conformations of gp41 are extended at the N-terminal ends of the helix, highlighted by the dashed box. Calculated all-atom root-mean-squared-deviation (RMSD) values against the HERV-K Env_Ecto_ complex or respective subunit are displayed under each structure.

In the HERV-K Env trimer, the TM-TM interaction is further reinforced by hydrogen bonds and salt bridges interfacing HR2 α10 with the adjacent TM fusion peptide and HR1 α6 (fig. 4g). This network ensures that the outer edges of the pre-fusion TMs are fastened together in the trimeric complex.

#### Glycosylation of Env

The HERV-K Env_Ecto_ sequence contains 10 possible N-glycosylation sites on each monomer, six on the SU and four on the TM (table S2). All 10 potential glycosylation sites are occupied and we were able to build the N-acetylglucosamine (NAG) cores for all the glycans (fig. S4a). For the glycan at N128, we could also model a mannose residue on each branch (fig. S4b). For the N566 glycan, we observe strong density corresponding to an α1-6 linked fucose (fig. S4c) which we modeled. Using the GlycoSHIELD pipeline(*64*), we modeled all glycosylation sites except for N566 as mannose-5. For each N566 site, we used a mannose-5 with an α1-6 linked fucose. Solvent-accessible surface area analysis of the glycosylated protein using a probe radius of 1.4 Å indicates that 31% of the surface of SU is shielded compared to the unglycosylated protein, with the apex particularly solvent-exposed. The TM is more shielded with 57% of its surface covered by glycans (fig. S4d,e). Together, the cryo-EM data indicate that HERV-K Env_Ecto_ has a 1:53 ratio of glycans to amino acids, and is less shielded than the envelope ectodomains of HIV-1 (1:23), SIV (1:27), and Jaagskiete Sheep Retrovirus (JSRV) (1:49), but more than Syncytin-1 from HERV-W (1:60) and Syncytin-2 from HERV-FRD (1:50)(*65–67*). Because HERV-K is typically silenced and integrated ubiquitously in all our cells, it is likely not subject to the same immune pressure as other exogenously circulating retroviruses.

### HERV-K Env adopts a novel fold

This structure of the HERV-K Envelope glycoprotein is the first pre-fusion trimer complex structure of any human endogenous retrovirus. The only other full endogenous retroviral Env structure yet available is from a hookworm endogenous viral element, a type II structure similar to flavivirus envelope proteins(*68*). Further, the HERV-K envelope structure presented here is also the first pre-fusion structure available for any betaretrovirus, whether endogenous or exogenous. Among all retroviruses, pre-fusion Env trimer structures are only thus far solved for the lentiviruses HIV-1 and SIV(*59*, *69*). For HERV-K Env, the only similarity in overall architecture is that in all three of HERV-K, HIV-1 and SIV Env, the SU domain is above (membrane distal) and the TM is below (membrane proximal), and that in all three retroviral trimers, the SU (or gp120) subunits are held in the pre-fusion trimer by the TM (or gp41), using a clasp or collar structure formed by the α helices from the HR1 and HR2 domains (fig. 3b,c). Beyond those two simple organizational similarities, the overall forms, secondary structural elements, and folds differ extensively between HERV-K Env and the HIV-1/SIV lentiviruses.

Comparison of the HIV-1 and SIV envelope trimer ectodomains thus far resolved to this structure of the HERV-K Env_Ecto_ reveals that HERV-K Env adopts a more streamlined form, primarily shaped by the SU domain. Compared to Env, the HIV and SIV trimers are shorter and wider, with heights ranging from 95 Å to 103 Å and widths between 108 Å to 120 Å (fig. 4a) vs. 117 Å tall and 83 Å wide for HERV-K Env.

The envelope glycoprotein of HIV-1 is highly dynamic, spontaneously transitioning between multiple conformational states which influence binding of antibodies, CD4, and co-receptors(*70*). Compared to each of the three described HIV-1 Env conformations, our structure of HERV-K Env_Ecto_ is more like the closed, unliganded conformation of HIV-1 gp140 (state-1; PDB:5CEZ), but the global root mean squared deviation (RMSD) is very high at 23 Å (fig. 4a). State-2 (PDB:7TFO), which is the occluded-open conformation triggered by CD4 binding, differs from HERV-K Env by an RMSD of 27 Å, and state-3 (PDB:5VN3), the CD4-bound, V3 loop exposed, open conformation, differs from this HERV-K Env structure by an RMSD of 38 Å. For HIV-1, conformational dynamics are required for receptor and coreceptor binding. In contrast, in HERV-K Env, the presumed receptor binding site is exposed at the trimer apex. During cryo-EM data processing, we did observe independent movement of each SU away from the vertical central axis, causing asymmetry (fig. S11a). The appearance of asymmetry in 3D classes suggests that some “breathing” flexibility of the SU domains may occur in HERV-K Env, potentially allowing adoption of more open or closed states.

The individual SU domain of HERV-K Env has a novel fold, as revealed by a systematic structural fold search utilizing Foldseek, DALI, and HHpred servers(*71–73*). The global RMSDs for HERV-K SU with any of HIV-1 gp120, SIV gp120, or Syncytin-2 SU are each >23 Å, and no other protein in the databases aligns at all with the HERV-K SU (fig. 4b). The area of greatest structural homology is the base domain of HERV-K, a region that is functionally analogous to the inner domain of HIV-1/SIV gp120. Both the HERV-K base domain and the HIV/SIV inner domain contain a distorted β barrel and a membrane-proximal β sheet (fig. 4b), and both are responsible for TM/gp41 interactions. Outside the base/inner domain, the SU structures are quite different. The HERV-K SU is vertically elongated and is composed primarily of β strands, with a discernable, 5-stranded β sheet in the core. In contrast, HIV-1 and SIV gp120s are a mixture of β sheets and α helices. Further, the lentiviral gp120s are larger than HERV-K SU (∼470-510 vs. 369 amino acids), and bear both an outer domain and V1V2 variable regions, while HERV-K is a single domain structure. The V1V2 loops of HIV/SIV are variable in sequence and critical for immune evasion. In contrast, the HERV-K sequence is highly conserved, likely a consequence of its endogenous nature and absence of immune-driven selective pressure (fig. S2). The conservation of all HERV-K Envs, however, is an advantage for the development of diagnostics and immunotherapeutics.

The human Syncytin-2 protein is critical for syncytiotrophoblast formation during placental development and is a co-opted envelope protein from the gammaretroviral HERV-FRD, inserted into the genome over 40 million years ago(*74*). Though the core domain of the Syncytin-2 SU subunit has been solved (PDB:7OIX), its full SU has not yet been resolved, either alone or in a trimeric complex with its TM. The Syncytin-2 SU core does, however, appear to share the β sheet formed by five antiparallel β strands extending upwards (fig. 4b). In Syncytin-2, the short loops that connect the antiparallel strands of this sheet form the binding site for MFSD2A, a membrane-embedded lysophosphatidylcholine lipid symporter(*75*), but it is not yet known whether the equivalent loops in HERV-K Env play a role in receptor or cofactor binding.

While the HERV-K SU is distinct from the lentiviruses, the TM_Ecto_ subunit in its prefusion conformation exhibits a fold that is more structurally similar to that of pre-fusion HIV-1 and SIV gp41, with global RMSDs ranging from 7-10 Å (fig. 4c). The TM/g41 of all three viral envelopes share similar tertiary structure, starting with the N-terminal fusion peptide that then becomes the C-shaped clasp predominantly composed of α helices which wrap around the SU/gp120 terminal β sheets. A long, vertically aligned α helix, which in HERV-K TM is α8 in the HR1 domain, is also present in HIV-1 and SIV gp41, and in each trimer, forms the central coiled-coil crucial for envelope trimerization (fig. 4c). This central coil structural feature is common among class I fusion proteins, even beyond retroviruses(*48*).

### The Post-Fusion Structure of HERV-K Env TM

Given the metastability of viral glycoproteins, both pre-fusion and post-fusion conformations of HERV-K Env are likely present on cells expressing Env. Thus, we sought to also solve the structure of the HERV-K TM ectodomain in its post-fusion conformation. The TM_Ecto_ (residues 498-603) was expressed with a maltose-binding protein (MBP) tag connected to TM_Ecto_ by a GS linker, a human rhinovirus (HRV) 3C protease cleavage site, and a T4 fibritin trimerization domain on the N terminus (fig. 5a). Residues 466-497 and 604-632 were not expressed due to the hydrophobicity of the fusion peptide and predicted flexibility. The purified TM_Ecto_ fusion protein was first complexed with Fab fragments of the Kenv-4 mAb, then treated with 3C protease to remove the MBP, which prevented aggregation of TM_Ecto_ alone (fig. 5a; fig. S5). The TM-Kenv-4 complex was purified by SEC and 696,758 particles from 19,225 movies were yield a 2.8 Å reconstruction by cryo-EM (fig. 5b; fig. S8). With the final map, we were able to model residues 502-548 and 567-603 (fig. 5c).

**Figure 5.**
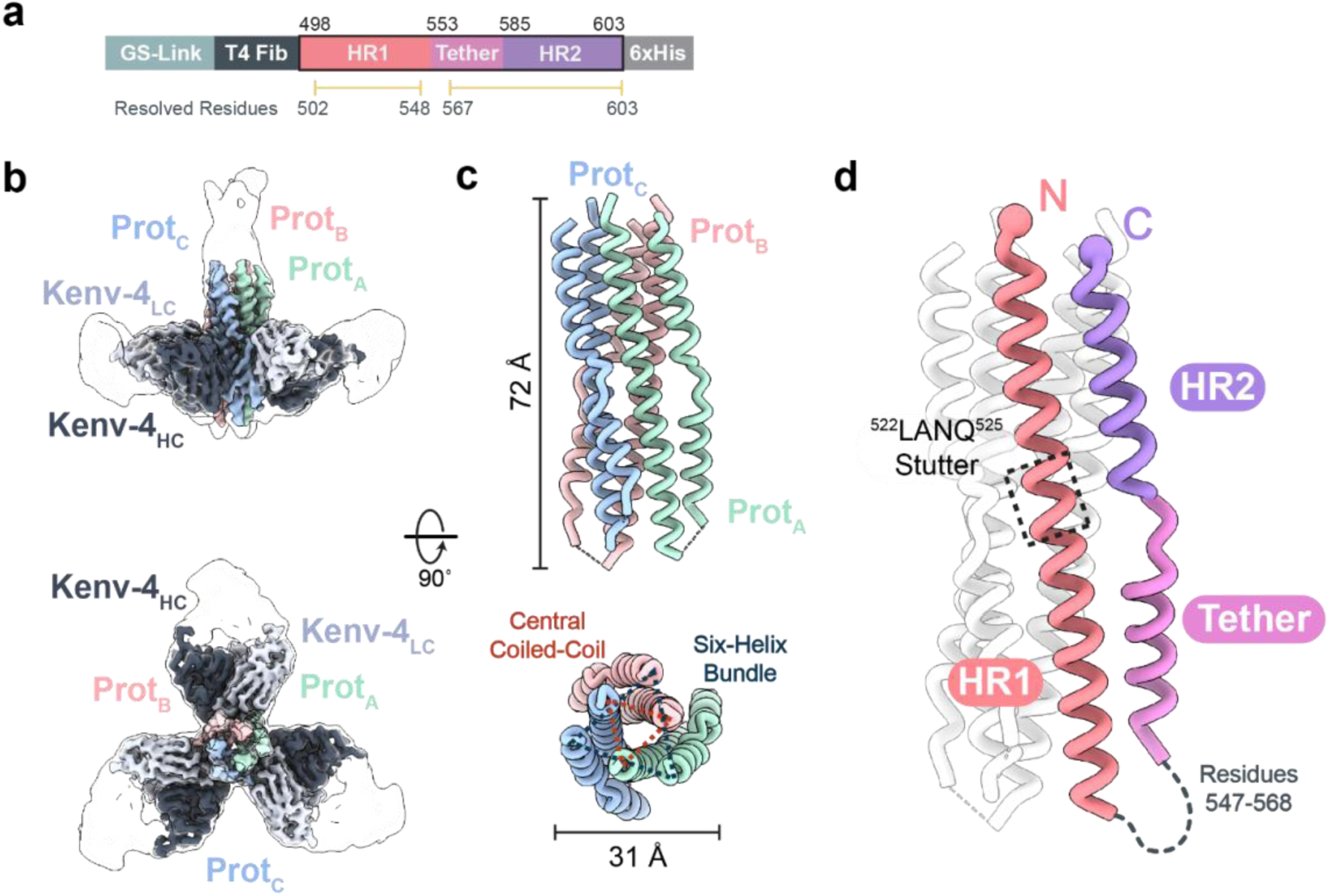
Cryo-EM structure of the Kenv-4 bound post-fusion HERV-K TM_Ecto_ complex. **a**. Schematic representation of the post-fusion HERV-K TM_Ecto_ construct. Abbreviations: GS-link, 17-amino acid linker; T4 fib, T4 fibritin trimerization domain; HR1, heptad repeat 1; HR2, Heptad repeat 2. **b**. 2.8 Å resolution cryo-EM map of post-fusion TM_Ecto_ trimer (colored using green, blue, and red shades) bound to three copies of the Kenv-4 Fab. A low-resolution map is overlaid and represented as a silhouette. **c**. A ribbon representation of the post-fusion TM_Ecto_. Six-helix bundle (6HB) and central coiled-coil structural features are shown using dotted triangles. **e**. A protomer of post-fusion TM_Ecto_ with domains colored as described in **a**.

The HERV-K TM ectodomain in the post-fusion conformation consists of heptad repeat 1 (HR1), an internal ‘tether’ region (includes the ^551^CX_7_C^559^ motif), and heptad repeat 2 (HR2) (fig. 5a,d). These three features assemble into a six-helix bundle (6HB), a structural hallmark of the post-fusion conformation in type I viral glycoproteins (fig. 5c,d). At the center of the 6HB, the HR1 helices form the central trimeric coiled coil, while the HR2 and tether helices adopt an antiparallel orientation, packing against the outer surface of the HR1 (fig. 5c).

The HR1 helix features the repeating seven amino-acid sequence pattern with hydrophobic residues occupying the a and d positions, facilitating inter-helical, hydrophobic packing interactions within the trimer (fig. S6). A four-residue stutter (^522^LANQ^525^) disrupts the regular heptad pattern in the HR1 region (fig. 5d), a feature missing from the HIV-1 and SIV lentivirus gp41, but observed in Influenza hemagglutinin (HA), Ebola Virus GP, Lymphocytic Choriomeningitis Virus GP, and endogenous retroviral envelopes: Syncytin-1 and Syncytin-2(*76–79*). This stutter causes deviations from the ideal helix packing geometry, leading to localized unwinding of the coiled-coil. This has been thought to impart flexibility to the helices and the trimeric core (*76*, *80*).

### The pre- to post-fusion transition

HERV-K Env-mediated fusion requires proteolytic processing of SU and TM, a low pH environment, and dissociation of SU from TM(*81*). The dissociation of SU from TM initiates the irreversible springing to the low energy state, where the fusion peptide inserts into the target membrane, starting the process of membrane fusion. Alignment of the pre-fusion TM subunit to the post-fusion TM revealed two major structural rearrangements that occur around α8, which remains largely unaltered between the two conformations. The first is the formation of the elongated HR1 helix (residues 502-546) by the displacement of the fusion peptide, pulling α6 and α7 away from the cell membrane, which are disjointed helices forming one arm of the C-shaped clamp in the pre-fusion conformation (fig. S7b). The HR1 central helix, initially ∼30 Å in length, extends to ∼67 Å. This is accompanied by the second rearrangement where the TM reverses direction. The α9 and α10 helices pack tightly along the now extended HR1 helix, forming the tether and HR2 helices, respectively, in the post-fusion structure (fig. S7c). This reorganization positions the sequence following HR2, including the unresolved membrane-spanning region at the C terminus, closer to the target membrane, driving membrane fusion (fig. S7c,d). Residues 546-566, which include an intramolecular disulfide bond between ^551^CX_7_C^559^ and β27, refold to facilitate the chain reversal. We could model this region in the pre-fusion structure, but could not resolve it in the post-fusion structure, likely due to inherent flexibility.

### Comparison to other retrovirus postfusion structures

We manually curated nine other retroviral envelope post-fusion structures from the Protein Data Bank (PDB) [HIV-1 (PDB:1I5X), SIV (PDB:1QBZ), ASLV (Avian sarcoma leukosis virus) (PDB:4JPR), HTLV-1 (Human T-lymphotropic virus 1) (PDB:1MG1), PERV (Porcine endogenous retrovirus) (PDB:7S94), Syncytin-1 (HERV-W) (PDB:6RX1), Syncytin-2 (HERV-FRD) (PDB:6RX3), XMRV (Xenotropic murine leukemia virus-related virus) (PDB:4JGS), MPMV (Mason-Pfizer monkey virus) (PDB:4JF3)] and performed sequence and structural alignments (fig. S8). All structures feature a long α helix corresponding to the HR1 region (fig. S8b,c). However, while other retroviruses possess only two helices (HR1 and HR2), the post-fusion HERV-K TM core exhibits an additional helix within the tether sequence, between HR1 and HR2, which we have termed the tether helix (fig. S8b,c). Interestingly, the HERV-K tether is closer in sequence to other retroviruses’ HR2 N termini, but it does not adopt the typical HR2 position (fig. S8b). Then HERV-K TM HR2 sequence aligns with the C termini of other HR2 sequences. Though we did not model the entire HERV-K HR2, the region that was modeled (residues 585-603) do pack similarly to the compared structures on the outside of HR1. RMSDs of the HERV-K TM with other retroviral TM (ranging from ∼4-20 Å for all atom pairs) are provided in table S3.

### Antibody binding interfaces

#### The Kenv-6 Epitope

Three Kenv-6 Fabs were solved in complex with the trimeric prefusion Env_Ecto_. Kenv-6 binds to a conformational epitope at the apex of the SU, and each copy of Kenv-6 buries 569 Å^2^ by the heavy chain and 363 Å^2^ by the light chain. The Kenv-6 heavy chain contacts Env primarily via the complementarity determining region (CDR)H3, which forms hydrogen bonds with both side chain and backbone atoms of residues in the loops of SU proceeding β13 (residues 277-281) and between β14-β15 (residues 309-313) and β17-β18 (residues 361-364) (fig. S9a). The CDRH1 and CDRH2 of Kenv-6 form hydrogen bonds to R280 and S281. Light chain CDRL1 and CDRL2 hydrogen bond to the hydroxyl oxygens on the sidechains of both Y367 and E314 and the backbone oxygen of R363 (fig. S9b). Notably, the Kenv-6 epitope is broadly conserved among intact HERV-K Envs encoded by the human genome (fig. S2), positioning Kenv-6 as a useful research tool for the detection of Env expression and folding. Indeed, we found that Kenv-6 was able to strongly stain neutrophils isolated from an SLE patient and an RA patient (fig. 1b). Neutrophils have been reported to express Envs from both type 1 (K102) and type 2 (K108) HERV-K insertions in these diseases (*30*, *31*).

#### The Kenv-4 Epitope

Three copies of the Kenv-4 Fab are symmetrically bound to the post-fusion HERV-K TM_Ecto_ trimer. The Kenv-4 epitope is located near the flexible loop (residues 581-585) of the TM, with each Fab spanning two adjacent monomers in their post-fusion conformation. The heavy chain of Kenv-4 buries 522 Å^2^ of surface area on the post fusion TM, while the light chain buries 386 Å^2^. The Kenv-4 heavy chain interacts with two adjacent monomers, while its light chain interacts with only one monomer. The heavy chain interaction is mediated by CDRH1 and CDRH2 (fig. S10a). Here, residue D32 of the CDRH1 salt bridges to R539 on one monomer (Prot_A_) and H571 on the adjoining monomer (Prot_B_); in the complex three copies of Kenv-4 make three such interactions around the trimer. Light chain binding is only mediated by CDRL1 with six hydrogen bonds, and Y70 in framework 3 hydrogen binding to E583 (fig. S10a). The post-fusion conformation-specific Kenv-4 did not react with neutrophils from either of the patient samples evaluated here, although other TM-binding mAbs did (fig. 1b).

## Discussion

HERVs have been present for millions of years and throughout human evolution. Although each one of us carries nearly 100 copies of the HERV-K (HML-2) provirus in our own genome, surprisingly, science thus far knows relatively little about the structures, functions and activities of these genes and their gene products when expressed. HERVs were previously thought to be silent “junk” DNA, but an emerging body of evidence now demonstrates activation of HERV elements during a variety of human disease states, including cancers and autoimmune diseases, and that patients of these diseases make antibodies against Env proteins. Among all HERV insertions, the HERV-K (HML-2) family stands out for its more recent integration into the human genome. As a result, many of the HERV-K insertions retain complete open reading frames and are able to express functional copies of the viral proteins(*4*, *6*, *10*).

Of the different HERV-K antigens, the envelope glycoprotein has been a particular focus as it is expressed on the surface of human cells and elicits human antibodies. Further, RNA transcripts and protein expression of Env have been demonstrated in various cancer tissues, including but not limited to breast cancer, ovarian cancer, leukemia, and melanoma(*13*, *15*, *19*, *20*, *82*, *83*). Expression of HERV-K Env is also associated with rheumatoid arthritis (RA), systemic lupus erythematosus (SLE), type I diabetes, multiple sclerosis, and various other autoimmune and neurodegenerative disorders(*9*, *28*, *31*, *84*, *85*).

The expressed Env has functions of its own: HERV-K Env binds to and co-immunoprecipitates with CD98HC (SLC3A2), an extracellular membrane-bound protein involved in amino acid transport, integrin signaling, and tumorigenesis. Binding of HERV-K Env to CD98HC activates the mTOR pathway, which regulates cell growth, proliferation, and tumor metabolism(*86*, *87*). Further, HERV-K Env expressed by the K108 locus on human chromosome 7 is a fully functional and fusogenic copy(*88*): expressed in multinucleated melanoma cells, HERV-K108 Env has been shown to mediate melanoma cell-cell fusion *in vitro*(*18*).

Expression of HERV-K Env proteins induces a humoral immune response against Env in cancer(*13*, *15*, *89*). Anti-Env autoantibodies are also elicited in various autoimmune disorders and may correlate with disease progression(*5*, *31*, *34–36*, *90*). Immune complexes formed between Env and these antibodies can activate immune cells and pro-inflammatory responses, contributing to the pathogenesis of diseases where anti-Env immunoglobins are prevalent (*5*).

Despite their appearance and apparent targetability in multiple human disease states, no structures of any HERV envelope trimer have been determined. Without such a template, we lack the means to understand how HERV-K Env interacts with cellular factors to trigger fusion or downstream cell signaling, how Env is recognized by human antibodies, if such antibodies can inhibit Env functions, how the immune response against Env may exacerbate disease pathology or confer protection, or how we may guide the design of targeted therapeutic interventions and Env-based diagnostics.

In this work, we raised a panel of novel monoclonal antibodies against HERV-K Env and used them to elucidate the first cryo-EM structures of the pre-fusion HERV-K Env (SU+TM) trimer ectodomain (2.2 Å resolution) and the post-fusion TM ectodomain (2.8 Å resolution). Though the conformations of Env on cell surfaces during disease states are not yet characterized, one may expect from our understanding of the metastability of viral glycoproteins that both pre- and post-fusion structures may exist on the cell surface and both be relevant for antibody recognition.

Kenv-6, which targets the folded SU subunit with preferential binding of the pre-fusion conformation, stains neutrophils from both RA and SLE. This indicates that SU is folding properly and is present in autoimmune disease-relevant cells. In these cells, the post-fusion-specific Kenv-4 did not stain the cells although other TM-specific antibodies did. Staining may be different in cancer cells than autoimmune cells. The tumor microenvironment is characterized by more acidic pH due to the accumulation of metabolic waste(*91*, *92*); recognition of the the acidic-pH-driven(*57*, *81*) post-fusion conformation of TM in Env-expressing cancers may be relevant in these contexts.

The overall structure of the pre-fusion HERV-K Env diverges from that of any retroviral envelope protein solved previously, with at least 23 Å RMSD from other known structures. The HERV-K Env trimer adopts a narrower profile than its lentiviral counterparts in HIV-1 and SIV and heavily incorporates β strands rather than α helices into the SU subunit. The 5-stranded β sheet at the core of the HERV-K Env SU, however, is also present in the SU core domain of Syncytin-2, a former endogenous retrovirus protein co-opted for human placental function(*75*). In the TM subunit of HERV-K, the core molecular machinery that guides 6HB formation and membrane fusion is similar to gp41 from HIV-1 and SIV. Other features, however, are unique, including a CX_7_C sequence motif found in betaretroviral envelopes(*46*) and the “tether” α helix in the post-fusion conformation. Further, HERV-K Env is not as glycosylated as exogenous retroviral envelopes (23 vs. 53 amino acids per glycan) and our analysis of its glycan shield reveals surfaces of vulnerability, particularly in SU, for targeting by anti-Env antibodies. These differences underscore the evolutionary divergence of HERV-K from other families of retroviruses.

The structures, stabilizing mutations, and antibodies presented here offer valuable templates and essential research tools that open doors for detection of HERV-K expression and analysis of its role in human disease states. HERV-K Env expression is observed in autoimmunity and five anti-Env antibodies described here stain human neutrophils from autoimmune disease patient samples, but not healthy neutrophils. In cancer, HERV-K Env has garnered significant interest as a possible neo-antigen or tumor-associated antigen(*21*, *93*). Indeed, strategies targeting Env for immunotherapy have been met with promising results: an anti-Env mAb termed 6H5 has been shown to inhibit growth and induce apoptosis of breast cancer cells *in vitro* in a dose-dependent manner. Used *in vivo*, mice bearing xenograft tumors displayed significantly reduced growth with 6H5 compared to an IgG control(*38*). Further, a nanobody targeting Env was reported to induce antibody-dependent cellular cytotoxicity in lymphoma cell lines and can be internalized by Env-expressing cells, warranting further exploration into anti-Env antibody-drug conjugates(*94*). Moreover, CAR-T cells engineered to target HERV-K Env have been effective in lysing breast cancer and melanoma tumors, inhibiting cell proliferation, and preventing metastasis *in vitro* and *in vivo*(*39, 40*). The GNK-301 mAb (GeNeuro) binds to a linear epitope located on the β14 strand at top of Env SU, blocks its interaction with CD98HC, and mitigates Env-related neurotoxicity in ALS(*33*). Our Kenv-6 mAb also binds to the SU, forming contacts with residues within the loop connecting β14 to β15 and the adjacent loop between β17-β18, although it is not yet known whether it has the same functionality.

Previous investigations to determine if there are autoantibody responses against Env in autoimmune diseases and immune responses in cancer have largely utilized linear peptides of Env or bacterially expressed Env, thereby failing to capture antibodies that target conformational epitopes or intact, glycosylated Env(*34*, *35*, *90*), and were performed in the absence of structural information on Env. The structures we describe now provide the blueprints for functional exploration of HERV-K Env. Further, the engineered and wild-type antigens described can be used in these studies, as well as to discover mAbs from patient samples and determination of which Env epitopes are immunodominant on the surface of cells. The results presented here also open doors for use of specific mAbs to detect or target diseased cells, or neutralize Env-mediated effects in disease, either by outcompeting patient-derived antibodies or by blocking binding to surface receptors.

This study establishes a structural basis for exploring the function(s) of the many HERV-K *env* loci and their gene products in disease pathogenesis, and provides tools needed for development of novel immunotherapies, diagnostics, and vaccines.

## Acknowledgements

The authors would like to thank S. Schendel for reviewing and editing the manuscript. We acknowledge the cryo-EM facility of La Jolla Institute for Immunology for assistance in grid preparation and data collection, and equipment supported by the GHR Foundation and other generous private donors.

## Author Contributions

J.S., K.M.H., and E.O.S. conceptualized and designed experiments. J.S., E.M.W., K.M.H., S.H., and P.P. contributed to the discovery, expression, and biochemical characterization of the antibodies. F.M. and T.M. collected and imaged neutrophils. J.S., E.M.W., C.Y., and V.I.L. designed, cloned, expressed, and purified the HERV-K Env_Ecto_ constructs. J.S., C.S., D.P., K.S., and R.R., cloned, expressed, and prepared post-fusion TM_Ecto_ proteins. J.S., C.S., and R.D.A. collected data for single particle analysis. J.S. performed data processing, model building, and refinement of the pre-fusion Env_Ecto_. C.S. performed data processing, model building, and refinement of the post-fusion TM_Ecto_. D.Z. helped with model refinement. J.S. and C.S. prepared the figures and wrote the original draft. K.M.H. and E.O.S. reviewed and edited the manuscript. All authors provided critical feedback and helped to shape the research, analysis, and writing of the paper.

## Data and materials availability

The Kenv-6 bound pre-fusion HERV-K Env_Ecto_ and Kenv-4 bound post-fusion TM_Ecto_ cryo-EM maps have been deposited to the EM Database with codes EMD-0000 and EMD-0000 respectively. Atomic models are deposited in the Protein Data Bank with accession codes 0ZZZ and 0ZZZ. After publication, all research materials referenced in this study will be made available upon reasonable request through material transfer agreements.

